# Model-based detection of putative synaptic connections from spike recordings with latency and type constraints

**DOI:** 10.1101/2020.02.12.944496

**Authors:** Naixin Ren, Shinya Ito, Hadi Hafizi, John M. Beggs, Ian H. Stevenson

**Affiliations:** Department of Psychological Sciences, University of Connecticut, Storrs, Connecticut, United States; Santa Cruz Institute for Particle Physics, University of California, Santa Cruz, Santa Cruz, California, United States; Department of Physics, Indiana University, Bloomington, Indiana, United States; Department of Biomedical Engineering, University of Connecticut, Storrs, Connecticut, United States

**Keywords:** synapses, spikes, multielectrode recording, Generalized Linear Model

## Abstract

Detecting synaptic connections using large-scale extracellular spike recordings presents a statistical challenge. While previous methods often treat the detection of each putative connection as a separate hypothesis test, here we develop a modeling approach that infers synaptic connections while incorporating circuit properties learned from the whole network. We use an extension of the Generalized Linear Model framework to describe the cross-correlograms between pairs of neurons and separate correlograms into two parts: a slowly varying effect due to background fluctuations and a fast, transient effect due to the synapse. We then use the observations from all putative connections in the recording to estimate two network properties: the presynaptic neuron type (excitatory or inhibitory) and the relationship between synaptic latency and distance between neurons. Constraining the presynaptic neuron’s type, synaptic latencies, and time constants improves synapse detection. In data from simulated networks, this model outperforms two previously developed synapse detection methods, especially on the weak connections. We also apply our model to *in vitro* multielectrode array recordings from mouse somatosensory cortex. Here our model automatically recovers plausible connections from hundreds of neurons, and the properties of the putative connections are largely consistent with previous research.

**New & Noteworthy:** Detecting synaptic connections using large-scale extracellular spike recordings is a difficult statistical problem. Here we develop an extension of a Generalized Linear Model that explicitly separates fast synaptic effects and slow background fluctuations in cross-correlograms between pairs of neurons while incorporating circuit properties learned from the whole network. This model outperforms two previously developed synapse detection methods in the simulated networks, and recovers plausible connections from hundreds of neurons in *in vitro* multielectrode array data.

## Introduction

Using *in vivo* or *in vitro* multielectrode arrays, the extracellular spiking of hundreds of neurons can be recorded simultaneously. These recordings are allowing new, large-scale studies of neuronal networks (Hahn et al. 2019; Harris et al. 2003; Levenstein et al. 2019; Okun et al. 2015; Tingley and Buzsáki 2018), and the number of neurons that can be simultaneously recorded is increasing approximately exponentially (Stevenson and Kording 2011). Depending on the species, brain area, and electrode configuration, these simultaneously recorded neurons can have tens of thousands of potential synapses between them. Detecting and characterizing these synapses represents a major challenge for neural data analysis. Here, we develop a model-based method incorporating network-level constraints on 1) the presynaptic neuron type and 2) the synaptic latencies between pre- and postsynaptic neurons. We examine whether these constraints can improve synapse detection using simulated data and large-scale *in vitro* multielectrode array recordings.

Detecting synaptic connections from extracellular spike observations is a difficult statistical problem. Since both spiking and synapses themselves are sparse, it is often difficult to distinguish between changes in spike probability that are due to a specific synaptic input, changes that are due other (typically unobserved) inputs, or due to chance. Using extracellular spike data, researchers often identify putative monosynaptic connections by examining cross-correlograms between the spiking of two neurons. If two neurons are connected, there will often be a fast-onset, short-latency peak (excitatory) or trough (inhibitory) in the cross-correlogram, where the post-synaptic neuron tends to spike more (excitatory) or less (inhibitory) following a pre-synaptic spike. Previous methods for distinguishing putative synaptic connections and non-connections in large-scale recordings used hypothesis testing to ask whether a peak or trough is significantly different from a baseline level of expected spiking (Barthó et al. 2004; Fetz et al. 1991; Fujisawa et al. 2008; Hatsopoulos et al. 2003; Perkel et al. 1967a). Recently, Kobayashi et al. (2019) developed a method that detects putative connections by applying a Generalized Linear Model to cross-correlograms (GLMCC), and used Matthews correlation coefficient (MCC, Matthews 1975) to find the optimal threshold for detecting connections. These methods typically treat decisions about the presence or absence of a synapse between each pair of neurons as separate hypothesis tests. However, synapses from the same presynaptic neuron are likely to share certain properties, and these shared properties could potentially improve the detection of synaptic connections. Here we aim to incorporate information from two basic features of neural circuits: 1) that neurons tend to be either excitatory or inhibitory and not both (Dale’s Law (Eccles et al. 1954)), and 2) that the synaptic latency between a pair of neurons should grow with the distance between the neurons (all else being equal). For example, knowing that there is an excitatory connection from neuron A to neuron B, increases the chances that other connections from neuron A should be excitatory. Similarly, if the distance between neuron A and B is known, then the latency of that connection provides some information about what latencies we might expect for neuron A’s other connections. These sources of information could potentially allow weak connections that are consistent with the circuit to be more readily detected and false positives due to noise to be rejected when that noise is inconsistent with the circuit.

To apply these circuit-level constraints, here we develop an extension of a Generalized Linear Model (extended GLM) to describe cross-correlograms between pairs of neurons and to automatically detect putative synaptic connections. Similar to (Kobayashi et al. 2019), here we fit an explicit model for the rate of post-synaptic spiking at each interval relative to the presynaptic neuron’s firing. This model includes both a fast, transient synaptic effect and a slower effect that accounts for potentially fluctuating baseline correlation. Here we add two constraints to the model: based on Dale’s law and the expected linear relationship between distance and synaptic latency, we rule out false positives by constraining presynaptic neuron type, synaptic latencies, and time constants. We then evaluate our model using two simulated integrate-and-fire networks. Our model outperforms previous synapse detection methods: spike jitter method and thresholding method, especially on the weak connections. We also apply our model to *in vitro* multielectrode array (MEA) data, where our model recovers plausible connections between hundreds of neurons in a slice culture of mouse somatosensory cortex. Many of the neurons appear to follow approximately linear distance-latency relationships, consistent with previous research *in vivo* (English et al. 2017), and neurons with excitatory/inhibitory connections often have waveforms that are wide/narrow, consistent with previous research *in vivo* (Barthó et al. 2004). Altogether, by incorporating constraints due to circuit structure, the model-based approach presented here may allow more accurate automated detection of synapses from large-scale spike recordings.

## Methods

### Extended Generalized Linear Model for Synaptic Detection

Here we develop an extension of a generalized linear model (extended GLM) to describe the spike correlograms between pairs of neurons: a suspected presynaptic neuron *i* and postsynaptic neuron *j*. For the binned spike trains of the two neurons, *n_i_* and *n_j_* (1 when there is a spike and 0 otherwise), the cross-correlogram is given by

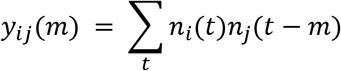

where *m* denotes the interval between pre- and postsynaptic spikes, and *y_ij_*(*m*) is the number of the times spikes in *n_i_* and *n_j_* are separated by an interval 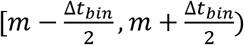 for bin size Δ*t_bin_*. Here in this study, we set the bin size Δ*t_bin_* to 0.5 ms, and model the intervals within a ±25ms window *m* = {−25, −24.5, ⋯, 25}*ms*

We then model the cross-correlogram using two components: 1) a slow effect caused by fluctuating firing rates and common input from other neurons, and 2) a fast effect caused by a potential synaptic connection. Namely, we model the rate of counts *λ_ij_* as a linear combination of the slow effect and the fast effect passed through an output nonlinearity:

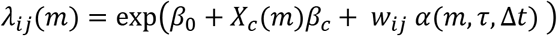

where *β*_0_ + *X_c_*(*m*)*β_c_* describes the slow effect and *w_ij_α*(*m*, *τ*, Δ*t*) describes the fast effect. For the slow effect, *X_c_*(*m*)represents a set of smooth basis functions learned by applying a low-rank, nonlinear matrix factorization to all the cross-correlograms in the dataset (see below). For the fast effect, we use an alpha function 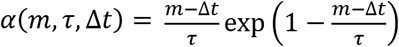 with a latency Δ*t* and a time constant *τ*, while *w_i,j_* represents the connection strength from neuron *i* to neuron *j* (positive for excitatory connections, negative for inhibitory). *β*_0_, *β_c_*, Δ*t*, *τ*, and *w_i,j_* are the parameters that are estimated in the model (see below for the details about optimization). Note that, since log *λ_ij_*(*m*) is nonlinear in the parameters *τ* and Δ*t*, this model is not a traditional GLM but an extension. The parameters, thus, cannot be optimized with traditional methods (e.g. iterative reweighted least squares).

In addition to this extended GLM (the full model), we also fit a reduced, slow model that is a GLM, *λ_ij_*(*m*) = exp(*β*_0_ + *X_c_*(*m*)*β_c_*), which only contains the basis functions without the alpha function to capture the synaptic effect. If the full model substantially outperforms the slow model, we can infer that there is putative synaptic connection from the pre- to postsynaptic neuron.

### Generating basis functions to describe the slow effect

To capture the slow fluctuations in correlograms, we use low-rank nonlinear matrix factorization to learn a set of smooth basis functions *X_c_*. Here we aim to reconstruct all of the correlograms in a given multielectrode recording using a generalized bilinear model:

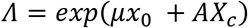

where *Λ* is a reconstruction matrix that aims to model the observed correlograms in terms of a vector of baseline correlations *μ*, a matrix of weights *A*, and the smooth basis functions *X_c_*. Note that here we model all *p* correlograms in the dataset simultaneously (*p* = *c*(*c* – 1)/2 if there are *c* neurons). To ensure that *X_c_* is smooth we further decompose this matrix as *X_c_* = *BX_s_* where *X_S_* is a set of cubic B-spline curves with equally spaced knots. Altogether, the matrix of correlograms is reconstructed using the parameters *μ*, *A* and *B*. *A* is a *p*×*n_β_* matrix, where *n_β_* is the number of basis functions that we aim to learn from the dataset (here set to 6). *B* is a *n_β_* × *n_s_* matrix, where *n_s_* is the number of spline curves (here set to 16). And *μ* is a vector that describes the baseline correlation for each correlogram, and that is multiplied by a row vector of ones *x*_0_. In order to estimate the parameters we use an alternating gradient descent algorithm to approximately maximize the overall log-likelihood 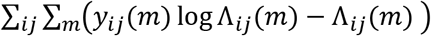. We alternate between updating the fits to each correlogram (*μ* and *A*) given a fixed set of bases (*B*) and updating the bases (*B*) given a fixed description of the individual correlograms (*μ* and *A*). Finally, we generate the basis functions as *X_c_* = *BX_S_*.

Although some pairs of neurons may have fast synaptic effects in addition to slower fluctuations due to common input, the proportion of these pairs is expected to be small. Since these connected pairs also have different weights, latencies, and time constants, the overall effect on the shapes of the learned bases *X_c_* should be relatively small.

### Parameter Estimation

We fit the cross-correlogram *y_ij_* using the full model 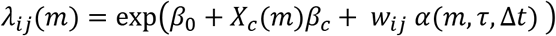 in two stages: 1) an initial fit that does not constrain the synaptic latencies, and 2) a subsequent fit that does. Using the estimated synaptic latency from the initial fit, we estimate the linear relationship between the distance and the synaptic latency. This enables us to use the estimated relationship to constrain the synaptic latency in the subsequent fit.

In stage 1, we fit the cross-correlogram *y_ij_* optimizing the penalized negative Poisson log-likelihood: 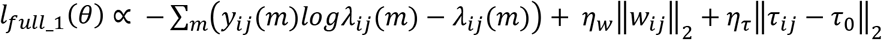, where 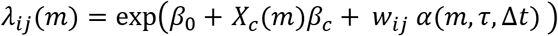 and estimate the parameters *θ* = {*β*_0_, *β_c_*, *w_ij_*, *τ*, Δ*t*}. This function is not convex due to the structure of the alpha function. However, we optimize the penalized log-likelihood using a non-linear conjugate gradient descent algorithm by using the minFunc toolbox in MATLAB (Schmidt 2005), and we use random restarts (50 times) in order to reduce the chances of getting stuck in local minima. Here *η_w_* and *η_τ_* are regularization hyperparameters that penalize large weights *w_ij_* and differences between the time constant *τ_ij_* from a reference *τ*_0_, respectively. Using the estimated latency Δ*t_ij_* from the initial fit and the distance between the neurons, we then estimate a “conduction” velocity *v_i_* and synaptic delay *dt_i_* for the presynaptic neuron *i* (see below).

In stage 2, using the estimated *v_i_* and *dt_i_*, we fit the cross-correlogram with an additional constraint on synaptic latency and estimate the parameters *θ* = {*β*_0_, *β_c_*, *w_ij_*, *τ*, Δ*t*}. Here we optimize the penalized negative Poisson log-likelihood:

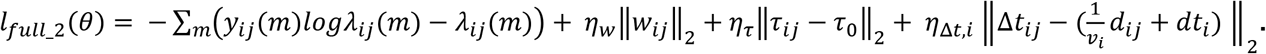

Adding the convex L2 penalty terms does not change the overall convexity of the function. Since the log-likelihood itself is not convex, here we again use a non-linear conjugate gradient descent algorithm with random restarts. *η_w_* and *η_τ_* are hyperparameters constraining the weight and time constant, as before. Given the distance between the two neurons *d_ij_*, the additional hyperparameter *η*_Δ*t,i*_ controls how strictly the synaptic latency Δ*t_ij_* should be tied to the predicted linear distance-latency relationship. Here *η_Δt,i_* is set based on the estimation of conduction velocity (see below, 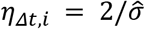).

In both stages, *w_ij_*, *τ_ij_*, *Δt_ij_* are log transformed so that they are strictly positive during the optimization (or, with a sign change, strictly negative when modeling an inhibitory *w_ij_*). In the results shown here we set *η_w_* = 5 and *η_τ_* = 20 through manual selection, and *τ_0_* is set to 0.8 ms. We have done a sensitivity analysis on the hyperparameters that we manually selected by doubling and halving the set values. We find that model performance is not very sensitive to the change in those hyperparameters, at least for the large-scale simulations we used here.

In addition to the full model, we also fit the cross-correlogram *y_ij_* using the slow model *λ_ij_*(*m*) = exp(*β*_0_ + *X_c_*(*m*)*β_c_*). We minimize the negative Poisson log-likelihood: 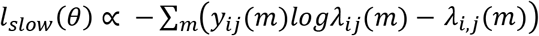 and estimate the parameters *θ* = {*β*_0_, *β_c_*} using iteratively reweighted least-squares. This provides a baseline null model, without a fast, synaptic effect.

After fitting, we compare the performance of the full model (stage 2) with the slow model by calculating the log likelihood ratio of the two models *LLR* = *l*_*full*_2_(*θ*) – *l_slow_*(*θ*). If the log likelihood ratio exceeds a certain threshold, we conclude that there is a putative connection from neuron *i* to neuron *j*.

### Structural constraints on fast, synaptic effects

While learned bases capture slow structure in the cross-correlograms across all pairs, we also aim to describe structure in the fast, synaptic effects for each presynaptic neuron. In the full model, we include two structural constraints: 1) we constrain the latency of synaptic connections to increase with increasing distance between neurons, and 2) we constrain presynaptic neurons to either excite or inhibit all of their postsynaptic targets, in accordance with Dale’s law. Together, these constraints have the potential to improve detection of weak connections that are consistent with the constraints and rule out the false positives that are inconsistent.

### Estimation of the “conduction velocity”

To implement the constraint that synaptic latencies should increase with distance, we estimate an approximate “conduction velocity” for each presynaptic neuron based on the distances between neurons and the estimated synaptic latencies from stage 1 above. Physiologically, conduction velocities vary as a function of axon diameter and myelination (Sakaguchi et al. 1993) so some differences are perhaps expected. However, that in most extracellular applications we are estimating the soma locations based on uncertain waveform information, and the locations of axons and dendrites are unknown. “Conduction velocity” is, thus, just an approximation of the potential positive relationship between synaptic latency and the distance.

Here we assume that there is a linear relationship between the synaptic latencies and the distances between the estimated somatic location of a presynaptic neuron *i* and postsynaptic neuron *j*,

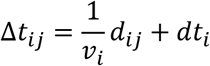

where Δ*t_ij_* is the synaptic latency, *d_ij_* is the distance between neurons, and the parameters *v_i_* and *dt_i_* describe the “conduction velocity” and “synaptic delay” of the presynaptic neuron. To estimate the parameters, we first fit all possible connections from the presynaptic neuron. Using initial estimates of 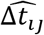 from the full model (stage 1), we then estimate *v_i_* and *dt_i_* for the neuron using a penalized weighted linear regression with the inter-neuronal distances as predictors. Namely, we minimize the penalized, weighted negative log-likelihood 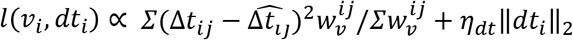, where the penalty ∥*dt_i_*∥_2_ ensures that *dt_i_* close to zero, and *η_dt_* is a hyperparameter, which we set to 5 based on manual search. The weights 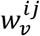 are set by ranking each pair of neurons based on the likelihood ratio between the slow model and full model in stage 1 (*l*_*full*_1_ – *l_slow_*, see **Parameter Estimation**), with the *r*th ranked pair having 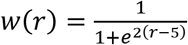. This allows the pairs that are more likely to be true connections (those with larger likelihood ratios) to have larger weights. Then, after conducting the penalized weighted linear regression, we pick the 5 neuron pairs with the largest weights to estimate the mean squared prediction error 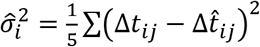, which measures the reliability of the estimation. This estimated prediction error, along with the estimated conduction velocity *v_i_* and delay *dt_i_*, is then used to constrain the penalized full model in stage 2 (see **Parameter Estimation** above).

### Estimation of the presynaptic neuron type

According to Dale’s Law, a single neuron should rarely be both excitatory and inhibitory, and connections with the same presynaptic neuron are most likely to be all excitatory or all inhibitory. In order to estimate the presynaptic neuron type, for each presynaptic neuron *i*, we fit all the cross-correlograms *y*_*i*1_, *y*_*i*2_, … *y_in_* using full model twice, once constraining *w_ij_* ≥ 0 (excitatory model) and once constraining *w_ij_* ≤ 0 (inhibitory model). Here we determine the presynaptic neuron type using the log likelihood ratio of the excitatory model fit to the inhibitory model fit.

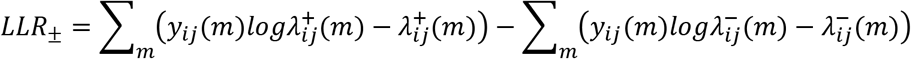

If the log likelihood ratio is positive, this suggests that the excitatory model provides a better description of the correlogram than the inhibitory model. For each presynaptic neuron, we use the single neuron pair with the largest likelihood ratio between two models to classify the neuron type (we tried using several weighting schemes, such as the average *LLR* across all pairs or the top-5 pairs, but for the simulations and datasets used here the top-1 pair performed the best of these schemes). We classify the presynaptic neuron *i* as a putative excitatory neuron if *LLR_±_* > 0, or as a putative inhibitory neuron if *LLR_±_* < 0. After the neuron type classification, we only adopt the corresponding full model (excitatory/inhibitory model based on the presynaptic neuron type) to later determine whether there is a putative synaptic connection. We label all the putative connections from an excitatory presynaptic neuron as putative excitatory connection, and all the putative connections from an inhibitory presynaptic neuron as putative inhibitory connections.

### Simulated networks of synaptically connected neurons

To examine how our model-based synapse detection approach performs we build two simulated networks of modified leaky integrate-and-fire (LIF) neurons. In real data, the shapes of cross-correlograms of two neurons can be affected by both the background activity of the network (external input shared by the network), and the patterns of presynaptic activity (e.g. high vs low firing rate, bursting). Here we designed two distinct simulations to capture these effects. In a first simulation we model a network of recurrently connected neurons that all receive background common input, creating slow fluctuations in the cross-correlograms similar to those observed in real data (Simulation 1 with common inputs). In a second simulation we then model a set of neurons receiving presynaptic inputs from experimentally observed spikes, creating presynaptic spike patterns similar to those present in real data (Simulation 2 with real presynaptic inputs).

For Simulation 1 with common inputs, we build a simplified, simulated network of adaptive leaky integrate-and-fire neurons with current-based synaptic inputs. 300 neurons are included in the simulation – 80% excitatory, 20% inhibitory. All the neurons are randomly distributed in a square area. The neurons are randomly connected, and only the neuron pairs whose distances are less than the median distance have synaptic connections. The connection probability is set to be 5% for the excitatory presynaptic neurons (Holmgren et al. 2003) and 20% for the inhibitory presynaptic neurons to balance the excitatory and inhibitory inputs (Barral and D’Reyes 2016; Dehghani et al. 2016; Haider et al. 2006). 60 minutes of current input and voltage recording for each neuron are simulated with a simulated sampling rate 10kHz for this network. The mean firing rate of all the neurons is 4.3Hz. In this modified LIF model (based on (Liu and Wang 2001)), the membrane potential dynamics are affected by three currents: 1) a leak current, 2) an after-hyperpolarization current, and 3) synaptic input

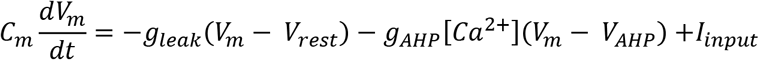

with

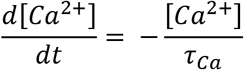

and if *V_m_*(*t*) = *V_th_* the neuron resets with

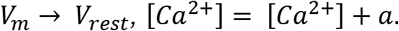

Here the dynamics of the membrane potential *V_m_* are governed by leaky integration of the input current, but every time the neuron spikes Ca-currents lead to an after-hyperpolarization, preventing the neuron from spiking rapidly. In the modified LIF model, when the membrane potential *V_m_* reaches the threshold *V_th_*, the neuron spikes, *V_m_* is reset to *V_rest_*, and [*Ca*^2+^] increases by the amount *a*.

The input current *I_input_* to each postsynaptic neuron is given by

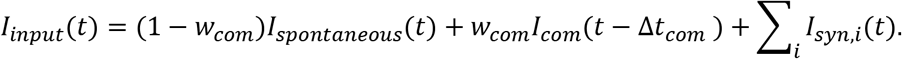

where *I_spontaneous_* is 1/*f* noise independently generated for each neuron, *I_com_* is 1/*f* noise shared by the whole network. Each neuron receives the common input with random latencies Δ*t_com_* to simulate the slow fluctuation caused by background common input, and *w_com_* is the random common input weight. *I_syn,i_* denotes the synaptic current from the *i*th presynaptic input added to the postsynaptic neuron with a synaptic latency Δ*t_ij_* after each presynaptic spike at *t_s_*, 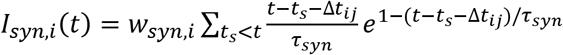. *w_syn,i_* is the synaptic weight randomly drawn from a bounded log-normal distribution – positive when the connection is excitatory and negative when the connection is inhibitory. Note that, since 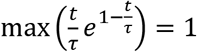, *w_syn_* sets the amplitude of individual Post synaptic current (PSC) in units of nA. Here we also give each presynaptic neuron a random “conduction velocity” *v_i_* and set the synaptic latency according to Δ*t_ij_* = *d_ij_*/*v_ij_*. This simulated network, thus, obeys the rule that synaptic latencies increase linearly with the distances between presynaptic neuron and postsynaptic neuron (see Table 1 for parameters).

**Table 1:**
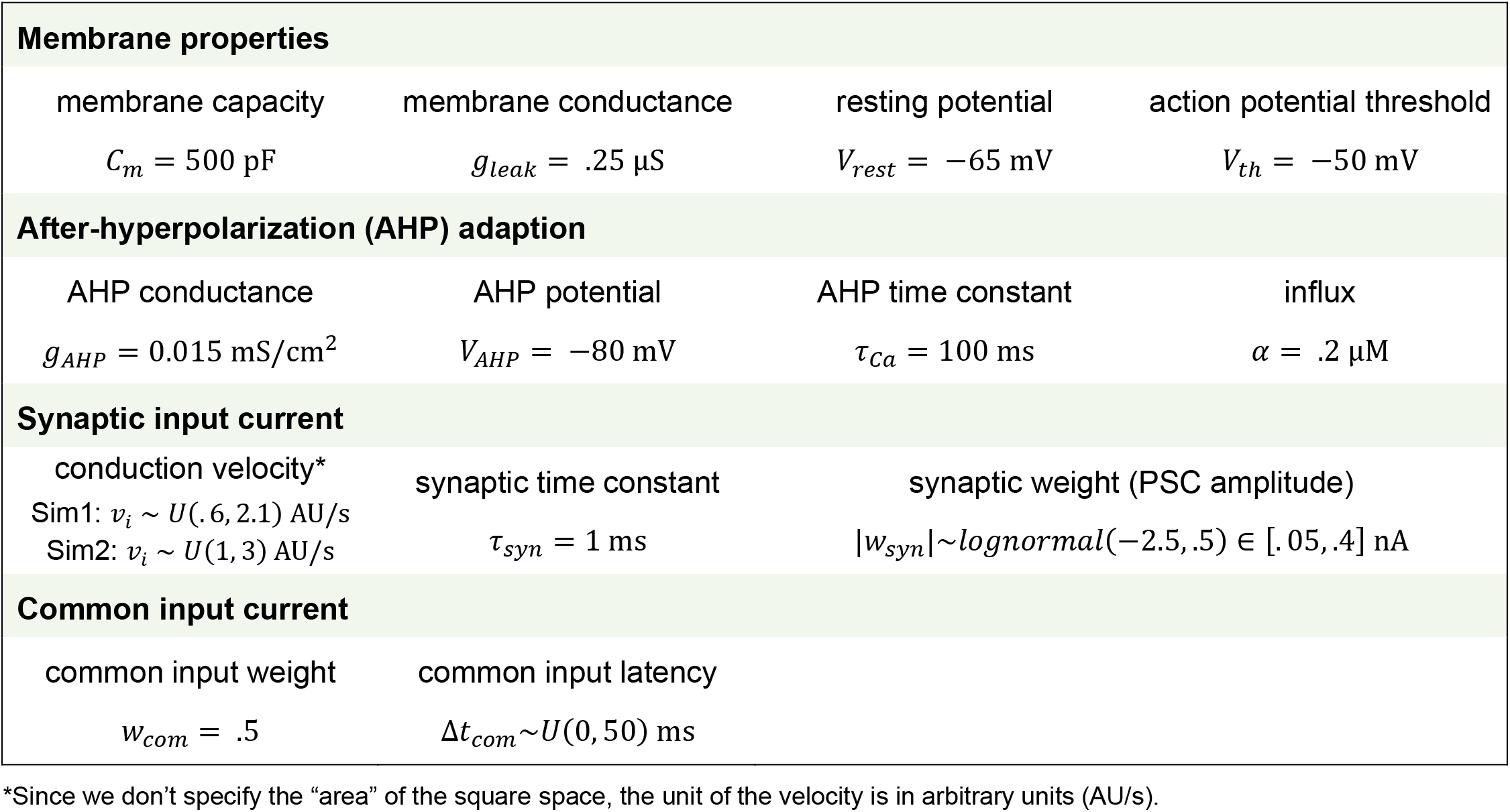
Parameters in the two simulated networks

In Simulation 2 with real presynaptic inputs, we model 300 adaptive leaky integrate-and-fire neurons that receive input from 300 neurons whose spike trains are from an *in vitro* multielectrode array recording. We randomly assign 80% of the 300 presynaptic neurons to be excitatory neurons and the rest to be inhibitory. The connection probability, connection rules, and LIF parameters are the same as in the first simulation (see Table 1). Here the simulated sampling rate is 20Hz, which was used in the *in vitro* recording, and the input currents do not contain the background common input, 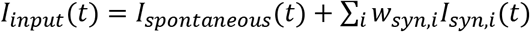.

### Synaptic detection based on hypothesis testing

In addition to our model-based synapse detection method we also examine two previous methods based on hypothesis testing: a thresholding method and a spike jitter method.

The thresholding method detects synapses by testing if the peak or trough in the correlogram is significantly different from the expected number of coincidences (Barthó et al. 2004; Perkel et al. 1967b). Here we model the count distribution using the mean 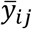 and standard deviation *s_ij_* of the cross-correlogram across bins – here between [−25,25] ms, excluding the bins within the interval of [-10,10] ms. We then compute the z-score 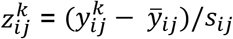 for each bin *k* and compare this to a critical value *z_c_*. If there is at least one bin within the interval of interest that exceeds the upper threshold *z_c_*, the connection from neuron *i* to neuron *j* is labeled as an excitatory connection. Similarly, if there is at least one bin within the interval below the lower threshold −*z_c_*, the connection from neuron *i* to neuron *j* is labeled as an inhibitory connection. In practice, the threshold *z_c_* can be adjusted to optimize the number of false positives/negatives. In comparing models, we use ROC curves to examine all thresholds (see below).

One potential problem with the thresholding method is that the baseline for a correlogram is often not constant. To address this, an alternative method (Fujisawa et al. 2008; Hatsopoulos et al. 2003) uses jittered spike trains to generate a baseline cross-correlogram that keeps the shape of the slow fluctuation while removing fast synaptic effects. With the jitter method, the presence of synaptic connections can then be inferred by testing if there is a peak or trough that is significantly different from this time-varying baseline. Here we use a variant of this method where, for each neuron, we randomly and independently jitter each spike on a uniform interval of [−5,5] ms (as in Fujisawa et al. 2008) and generate 1000 jittered spike trains. The baseline cross-correlogram between neurons *i* and *j* is then defined as the mean of the 1000 cross-correlograms constructed using the original spike trains of neuron *i* and the 1000 jittered spike trains of neuron *j*. We calculate the mean 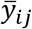 and standard deviation *s_ij_* of the 1000 cross-correlograms for each neuron pair. We then compute the z-score of each bin based on the original correlogram 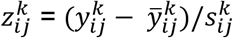. As in the thresholding method, if at least one of the bins within the interval of [0,10] ms exceeds the upper threshold *z_c_*, the connection is labeled as excitatory. Similarly, if there is at least one bin within the interval below the lower threshold −*z_c_*, the connection is labeled inhibitory.

### Evaluating methods for synapse detection

Using the simulations described above we evaluate our model-based synapse detection method alongside the thresholding method and jitter method. Benchmarking the performance of synapse detection methods on real extracellular recordings is difficult,since we are almost always uncertain about whether or not two neurons are monosynaptically connected. However, with simulations, the ground-truth connectivity is known, and we can compare the detection accuracy for different methods. Here we use receiver operating characteristic (ROC) curves, specifically comparing false positive and true positive rates. Since the number of true positives is small (less that ~5%), these rates and the area under the ROC curve (AUC) give a more accurate impression of the detection performance than the overall accuracy and can be calculated without a set threshold. The scores we use to determine whether there is a synaptic connection in generating the ROC curves vary for the three methods. For the model-based method developed here, we use the log likelihood ratios of full model to slow model, while for thresholding and jitter methods, we use the largest z-score within the [0,10] ms interval.

The ROC curves measure the overall performance of different methods on a series of thresholds. But when we apply the method to real data and plan to make decisions on synapse detection, we still need to specify a threshold. The choice of threshold has a large effect on the detection of putative synaptic connections. A threshold that is too strict will result in a large number of false negatives, while a threshold that is not strict enough will result in a large number of false positives. The uncertainty and diversity of the real datasets make it difficult to pick the optimal threshold. Here, for illustration, we pick the threshold based on the results in our simulated network (we pick Simulation 1 here since the threshold based on Simulation 1 is stricter). Since synaptic connections are relatively rare compared to the total number of neuron pairs, we use Matthews correlation coefficient (MCC, Matthews 1975) to measure the performance of different thresholds, which performs well for imbalanced data (Boughorbel et al. 2017):

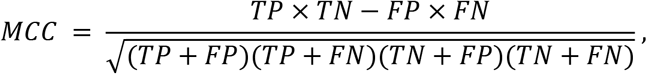

where *TP* is the number of true positives, *TN* is the number of true negatives, *FP* is the number of false negatives, and *FN* is the number of false negatives. For the model-based method, the maximum MCC is .86 (TPR = 84.83%, FPR = 0.09%) for Simulation 1, the corresponding threshold is 4.64 (log likelihood ratio). It may be valuable to note that this threshold is relatively close to the decision rule that would be given by minimizing the Akaike or Bayesian Information Criteria (AIC or BIC), where the log likelihood ratios would need to be greater than 3 or ~6.9, respectively (based on *k* = 3 extra parameters and *n* = 100 bins of observations). For jitter method, the maximum MCC is .69 (TPR = 60.23%, FPR = 1.02%), the corresponding threshold is 3.59 (z-score) for Simulation 1. In comparing the results from different synapse detection methods with real data, we pick the thresholds for our method and the jitter method based on these maximum MCC results from the simulation.

In addition to the choice of threshold, the jitter method has 1 hyperparameter (jitter interval) and the model-based method has 7 (*η_w_,η_τ_,τ_0_,η_Δt,i_,η_dt_,n_β_,n_s_*) that are used for the entire set of putative connections. Here we fix the hyperparameters for the model-based approach based on a coarse, manual optimization that minimizes false positive fits with unlikely latencies (Δ*t*) and time constants (*τ*). These values could also potentially be optimized using the cross-validated likelihood but, in practice, the results are robust across a wide range of settings.

### MEA data

To examine how these methods detect putative synaptic connections in experimental data we use *in vitro* recordings of spontaneous activity from organotypic slice cultures of mouse somatosensory cortex made using a large and dense multielectrode array (512 electrodes, 60 μm interelectrode spacing, 5 μm electrode diameter, flat electrodes, roughly 1 mm by 2 mm total array area). The angle of the slices was approximately coronal, but the lateral side of the plane was advanced by 15 degrees in the anterior direction. The extracellular signals were recorded for 60 minutes at 20 kHz, and the spiking activity was then spike sorted based on the waveforms of the marked electrode and its six adjacent neighbors using principal component analysis (PCA). The location of each neuron was estimated using a 2D Gaussian fit to the maximum values of the spike triggered average waveforms across multiple electrodes. There are 25 datasets available, most of which possess hundreds of neurons (min: 98, max: 594, mean: 309, total: 7735, mean firing rate of the neurons: 2.1 Hz). All data is available via the Collaborative Research in Computational Neuroscience (CRNCS) Data Sharing Initiative: https://crcns.org/data-sets/ssc/ssc-3/about-ssc-3. Additional experimental details and raster plots can be found in (Ito et al. 2014, Fig. 4A).

To simulate the network with real data input (Simulation 2), we used spike trains from the highest firing rate neurons combined from two datasets (datasets 16 and 23), choosing 300 neurons in total (out of 904 possible). The mean firing rate of the 300 neurons was 5.57Hz (min: 1.88Hz, max: 44.55Hz).

For examining putative synaptic connectivity in the experimental data, we use dataset #13 (number of neurons: 381, mean firing rate: 1.95 Hz) and dataset #23 (number of neurons: 310, mean firing rate: 2.81 Hz). Here we exclude the neurons with less than 1000 spikes recorded, 68 neurons (17.85%) are excluded from dataset #13, 21 neurons (6.77%) are excluded from dataset #23. Before we apply the detection methods on these datasets, we also exclude the neuron pairs where the correlogram may be misestimated due to the way that spike trains were sorted. If the waveforms of two neurons show up on the same set of electrodes, near simultaneous spikes tend to overlap and be sorted inaccurately (Pillow et al. 2013, “spike shadowing”). Here, we calculate a spike sorting index 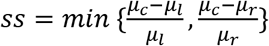 to exclude the cross-correlograms with a peak or trough near *m* = 0. Here *μ_c_* is the total number of counts within 1.5 ms of the center of the correlogram (3 bins), *μ_l_* is the total number of counts within 1.5 ms (3 bins) that are to the left of the center, *μ_r_* is the total number of counts within 1.5 ms (3 bins) that are to the right of the center. We exclude the neuron pairs when the spike sorting index *ss* is greater than 0.5. Based on this rule, 6.51% of the neuron pairs are excluded from dataset #13, 5.55% of the neuron pairs are excluded from dataset #23.

## Results

Here we develop an extension of a generalized linear model (GLM) to describe the correlograms between pairs of neurons. This model aims to separate the cross-correlogram between each pair of neurons into two parts: 1) a slow effect caused by fluctuating firing rates and common input from other neurons, which is fit using a group of smooth basis functions learned from the data, and 2) a fast effect caused by the synaptic connection, which is fit by a short-latency, fast onset alpha function (Fig. 1A). In this study, we model the time interval between −25 ms to 25 ms, with a binsize of 0.5 ms. To determine whether or not a given pair of neurons might be synaptically connected we then compare the full model with a reduced model that has the slow effect but not the fast effect. If the full model provides a better description of the data than the slow model (using log-likelihood ratio), this may indicate that there is a synaptic connection between the two neurons (Fig. 1B).

**Figure 1:**
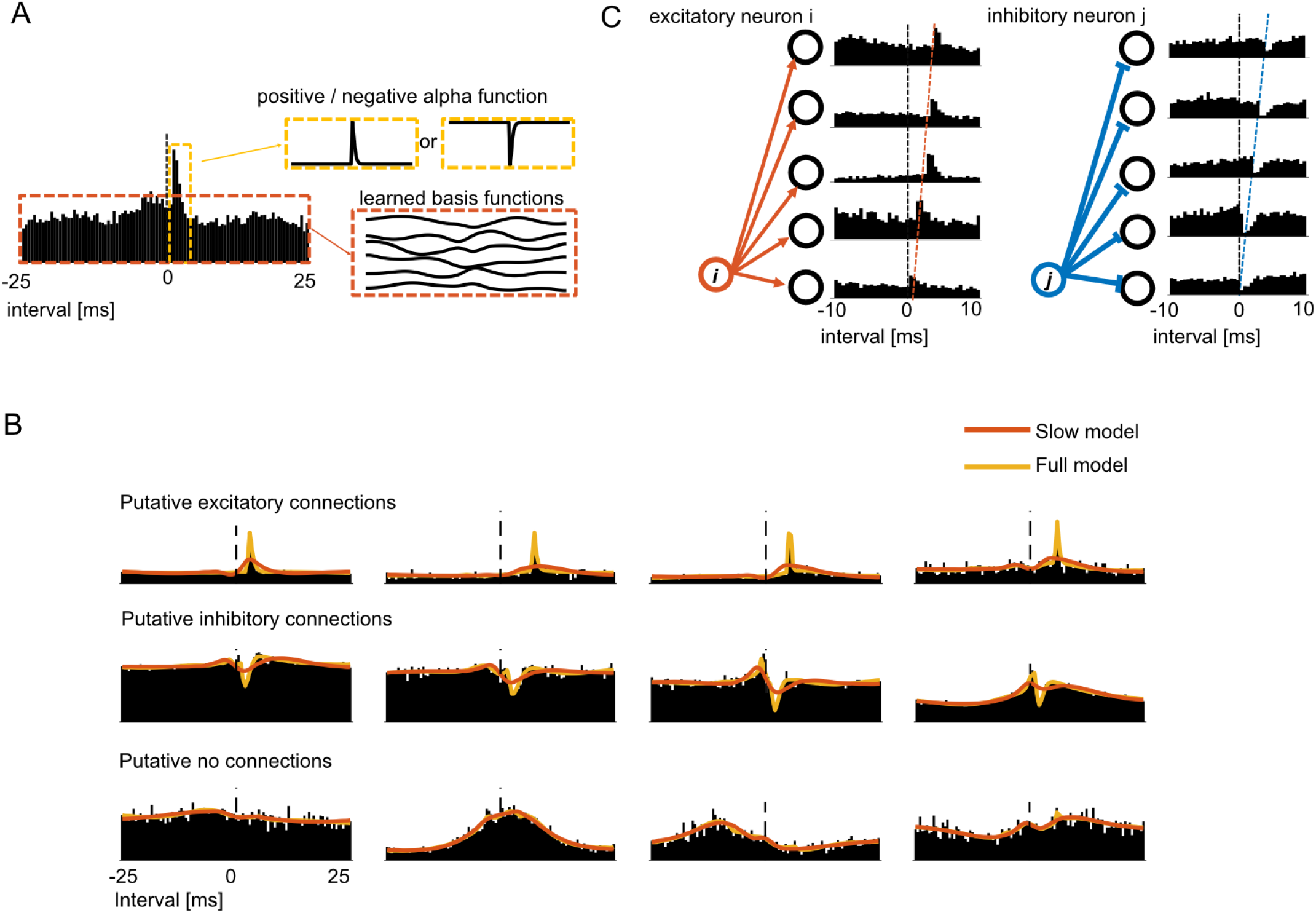
Model-based description of the cross-correlogram between the spiking of a pair of potentially connected neurons. A: The extended GLM separates the cross-correlogram into two parts: 1) a slow effect that we fit using a group of smooth basis functions which were learned from the whole network (outlined in red), and 2) a fast effect that we fit using a short-latency, fast onset alpha function (outlined in yellow). B: Some examples of model fits for cross-correlograms of putative excitatory, putative inhibitory, and putative non-connections. If the full model (yellow) provides a better fit to the correlogram than the slow model (red), we label the neuron pair as a putative connection. C: The schematic figure shows the structural information that we use to constrain the model fits: 1) the connections from one presynaptic neuron should be either all excitatory or inhibitory, and 2) the synaptic latency should increase with increasing distance between pre- and postsynaptic neurons.

Although this model comparison based on the correlogram between a single isolated pair of neurons can provide evidence of a putative synaptic connection, incorporating information from other connections may be able to improve detection accuracy. Here we first constrain the parameters of the full model based on the presynaptic neuron type. Since neurons are rarely both excitatory and inhibitory (Dale’s Law), synaptic connections with the same presynaptic neuron are most likely to be all of one sign. If a presynaptic neuron has a connection with a clear positive synaptic effect, this can indicate that other connections from this presynaptic neuron should be positive as well. Second, we constrain the parameters of the full model based on the synaptic latency. Synaptic latencies tend to increase with the distance between the pre- and postsynaptic neuron (Fig. 1C). Here we assume a linear relationship between distance and latency and estimate a “conduction velocity” for each presynaptic neuron. If this relationship is clearly linear, the possible latencies for other connections can be constrained. Together, these two constraints may act to better detect the weak connections and exclude the false positives that violate the expected structure (see more details in methods).

### Simulated networks with type and latency constraints

To evaluate our model, we build two simulated networks of adaptive leaky integrate-and-fire (LIF) neurons: Simulation 1 with common inputs, a network of 300 recurrently connected LIF neurons receiving slow, background common input, and Simulation 2 with real presynaptic inputs, a network of 300 unconnected LIF neurons receiving input from a set of experimentally recorded spike trains. In the first simulation, 80% of the neurons are randomly assigned to be excitatory, with the rest being inhibitory. The neurons are randomly connected to each other with a connection probability of 5% for the excitatory presynaptic neurons and 20% for the inhibitory presynaptic neurons. Synaptic weights (as PSC amplitude) are then randomly drawn from a log-normal distribution, similar to results from *in vitro* observations (Song et al. 2005). In addition to the synaptic input, all neurons receive background common input from a single slowly fluctuating, noisy source with a random delay (see Methods). This common input produces baseline fluctuations in the cross-correlograms similar to what is frequently observed in the real data (Fig 2A). Additionally, we assign each presynaptic neuron a “conduction velocity” and make the synaptic latencies between neurons distance-dependent. In Simulation 1, the mean firing rate of all the neurons is 4.35 Hz (min: 2.58 Hz, Q1: 3.72Hz, Q2:4.23 Hz, Q3: 4.78 Hz, max: 8.71 Hz, SD = .89 Hz). This simulated network, thus, has realistic slow fluctuations in the correlograms, obeys Dale’s Law, and the relationship between synaptic latency and distance increases linearly for each presynaptic neuron.

**Figure 2:**
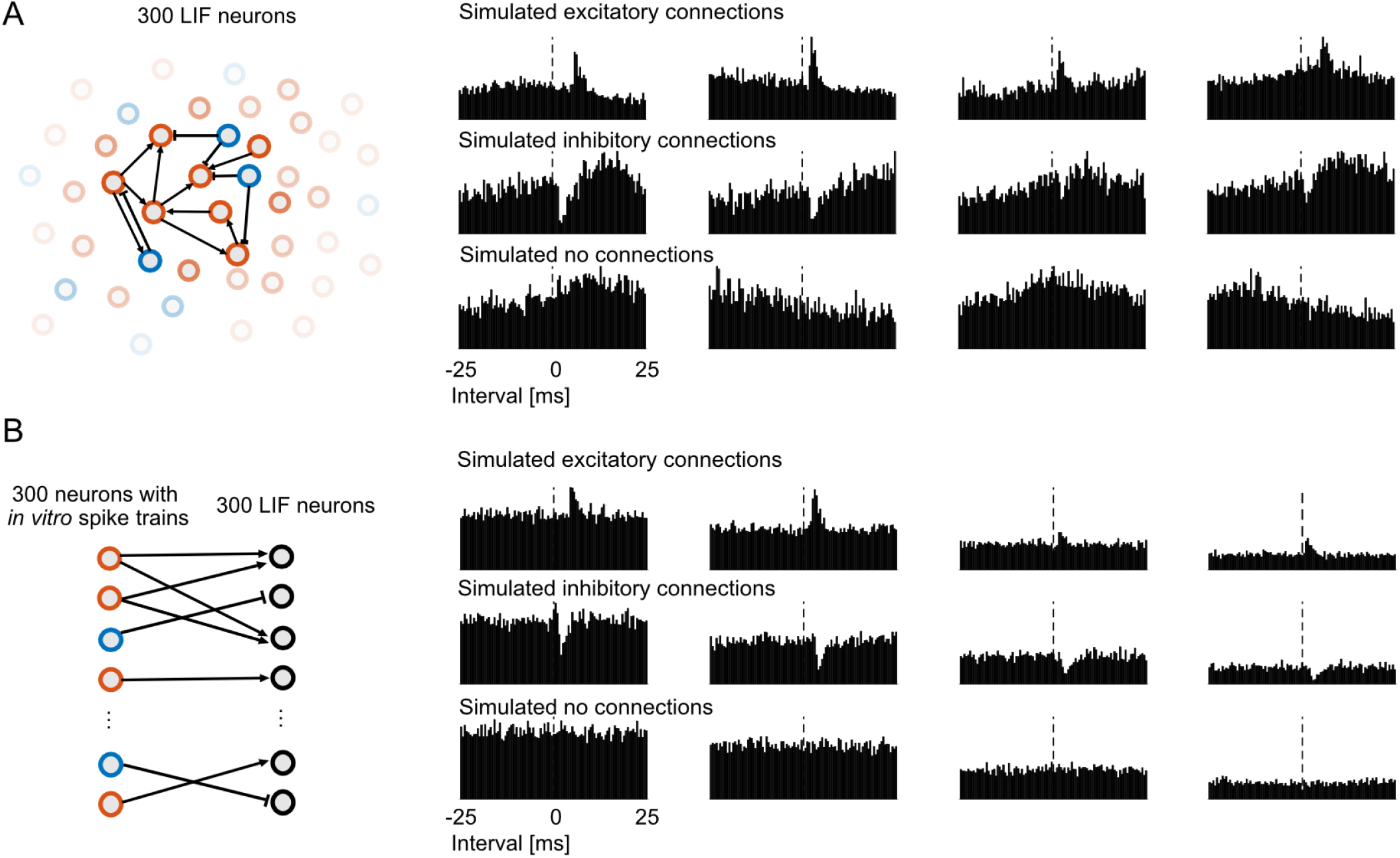
Two simulated networks of leaky integrate-and-fire (LIF) neurons. A: Schematic showing the structure of Simulation 1 with common inputs (left). 300 LIF neurons (80% excitatory, 20% inhibitory) are randomly connected to each other with constraints on synaptic latency (see Methods). They receive background common input to generate a slow baseline fluctuation in the cross-correlogram. Examples of the cross-correlograms for simulated excitatory, inhibitory and non-connections (right). B: Schematic showing the structure of Simulation 2 with real presynaptic inputs (left). 300 LIF neurons receive presynaptic inputs from 300 experimentally recorded spike trains. We randomly assign 80% of the presynaptic neurons to be excitatory and the rest to be inhibitory. Note that, although the schematic illustrates the bipartite connectivity structure, the 600 neurons are randomly distributed in space and the synaptic latencies increase linearly with distance between the neurons as in Simulation 1. Examples of the cross-correlograms of simulated excitatory, inhibitory, and non-connections from the second simulation (right). Due to the fact that the experimentally recorded spike trains have greater variation in the average firing rates and patterns, the cross-correlograms here have a wider range of absolute baselines.

The second simulation consists of a set of 300 LIF neurons each receiving presynaptic inputs from a subset of 300 spike trains recorded *in vitro*. Again, the presynaptic neurons are randomly assigned to be excitatory (80%) or inhibitory (20%). The presynaptic neurons are randomly connected to the postsynaptic neurons with a connection probability of 5% for the excitatory presynaptic neurons and 20% for the inhibitory presynaptic neurons, and the synaptic weights are randomly drawn from a log-normal distribution. The synaptic latencies also increase linearly with distance, as before. In this case, although there is no common input, the presynaptic spike patterns are drawn from experimental recordings and the presynaptic neurons have greater variation in their firings rates and inter-spike interval patterns. The mean firing rate of the presynaptic neurons in this simulation is 5.57 Hz (min: 1.88 Hz, Q1 = 2.84 Hz, Q2 = 4.26 Hz, Q3 = 6.55 Hz, max: 44.6 Hz, SD =4.98 Hz). The mean firing rate of the postsynaptic, LIF neurons is 5.63 Hz (min: 4.00 Hz, Q1: 5.10 Hz, Q2: 5.56 Hz, Q3: 6.10, max: 8.89 Hz, SD = .80 Hz). Although the correlograms of Simulation 2 do not have slow baseline fluctuations (Fig 2B), they have a broader range of absolute baselines and will allow us to determine to what extent synapse detection is affected by more realistic presynaptic spike patterns.

A central assumption of the model-based detection approach used here is that neuron type and latency constraints can, in principle, allow information to be shared across the connections made by a presynaptic neuron. However, in order for these constraints to be useful, the model must be able to accurately estimate both whether a presynaptic neuron is excitatory or inhibitory and the presynaptic neuron’s “conduction velocity” from noisy spiking data. Therefore, before evaluating whether these constraints improve detection, we determine how accurately we can recover neuron type and “conduction velocity” in each of the simulations.

In order to determine the presynaptic neuron type, we compare two models of the cross-correlogram between each pair of neurons: one with a positive fast, synaptic effect and the other with a negative synaptic effect. We can then estimate the type of each presynaptic neuron by asking which of the two models provides a better description of the cross-correlograms involving that presynaptic neuron (see Methods). Using our model, in Simulation 1 with common inputs, 97.7% of the neurons are labeled correctly (7 out of 300 mislabeled). In Simulation 2 with real presynaptic inputs, 95% of the neurons are labeled correctly (15 out of 300 mislabeled). In this case the mislabeled neurons are also relatively low-firing rate (mean firing rate = 3.31 Hz, compared to 5.57 Hz for all presynaptic neurons).

We then evaluate how well we can estimate each presynaptic neuron’s conduction velocity from the cross-correlograms. Here we estimate the synaptic latency between each pair of neurons and use a weighted linear regression to then estimate the “conduction velocity” of each presynaptic neuron (see Methods). Using this approach, we find that we can recover the true velocity that was assigned to each of the presynaptic neurons in the simulations relatively accurately. For Simulation 1 with common inputs, the estimated latency-distance parameters are correlated with their true values 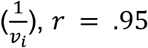, *p* < .01, root mean squared error *RMSE* = .0013 AU/s (Fig. 3A) and for Simulation 2 with real presynaptic inputs, *r* = .67, *p* < .01, *RMSE* = .0015 AU/s (Fig. 3D).

**Figure 3:**
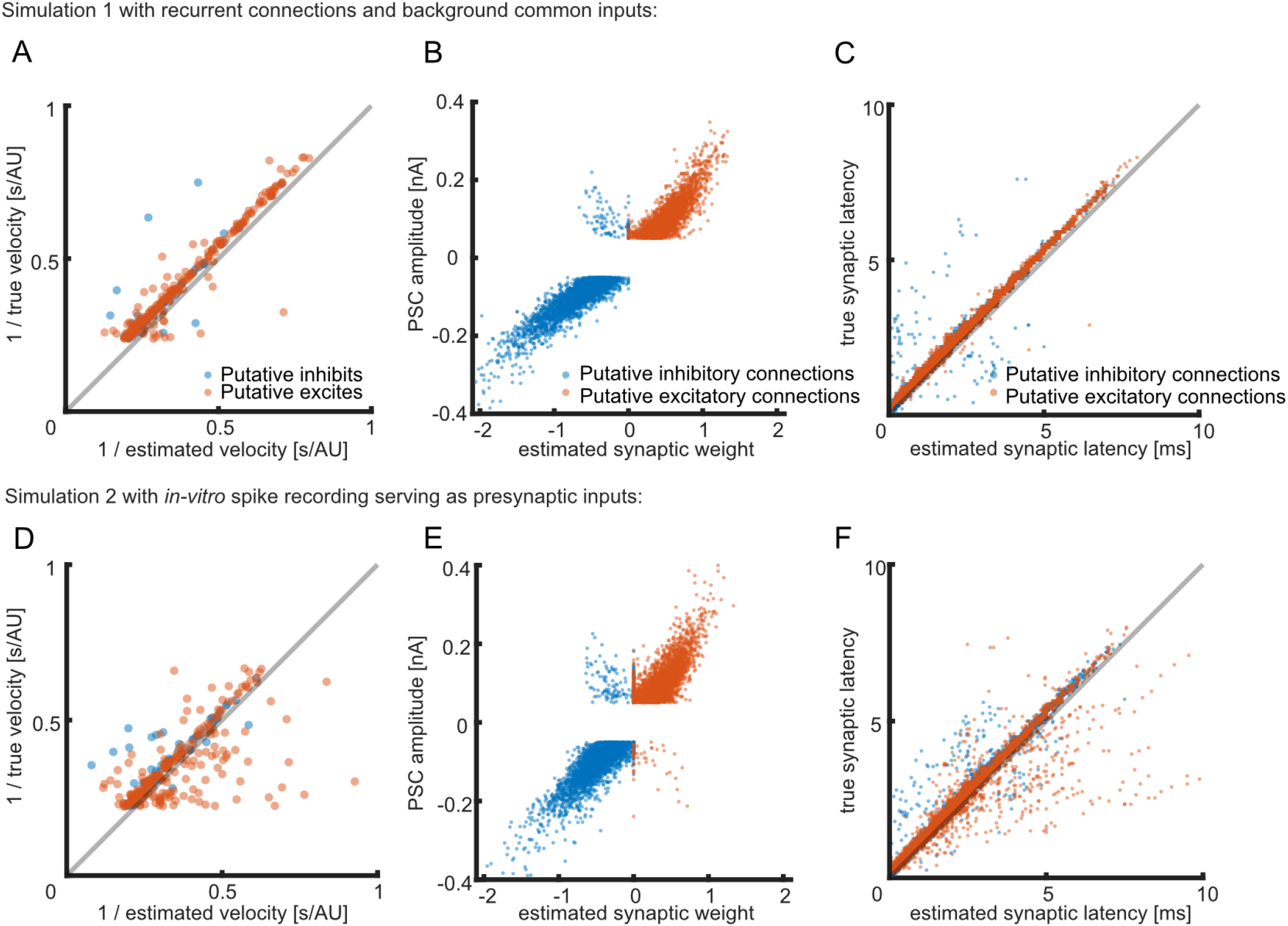
The extended GLM can capture the properties of presynaptic neurons and individual synaptic connections in two simulated networks: a recurrent network with common input (A-C) and a network with realistic input (D-F). A & D: Estimated and true presynaptic conduction velocity. Each dot represents one simulated presynaptic neuron. Colors indicate the estimated presynaptic neuron type. Axes are normalized. B & E: Estimated and simulated synaptic weight (*w_ij_*, coefficient of the alpha function). Here each dot represents one true connection. Y-axis is the PSC amplitude assigned in the simulations. Note that dots in the second and fourth quadrants correspond to cases where the presynaptic neuron type has been misestimated. C & F: Estimated and simulated synaptic latency. Again, each dot represents one true connection.

Given these constraints, we can then examine how well we are able to recover the properties of individual connections. Here we analyze only the true connections within the simulations and find that the true synaptic weight can be recovered relatively accurately: for Simulation 1 with common inputs, *r* = .96, *p* < .01, for Simulation 2 with real presynaptic inputs, *r* = .92, *p* < .01. Similarly, synaptic latency can be estimated accurately: for Simulation 1 with common inputs, *r* = .98, *p* < .01, *RMSE* = .37 ms, for Simulation 2 with real presynaptic inputs, *r* = .93, *p* < .01, *RMSE* = .58 ms. And Including neuron type and latency constraints improves those reconstructions (For the model without constraints, Simulation 1: latency: *r* = .73, *p* < .01, *RMSE* = 1.16 ms, weight: *r* = .92, *p* < .01. Simulation 2: latency: *r* = .38, *p* < .01, *RMSE* = 2.32 ms, weight: *r* = .79, *p* < .01). Together, these results illustrate how, for simulated networks, our model is able to capture the type and conduction velocity of presynaptic neurons, as well as the parameters of individual connections.

### Synapse detection with simulated spike trains: Evaluating the model-based method

Given that the model-based approach can recover the properties of presynaptic neurons (type and conduction velocity) and the properties of individual connections, we then ask how well our model can distinguish which pairs of simulated neurons are synaptically connected and which are not. We applied our model and two previously used synapse detection methods: the thresholding method and spike jitter method, to the two simulations described above. Briefly, the thresholding method is based on testing if the peak or trough in the correlogram immediately following a presynaptic spike is significantly different from a constant, baseline number of coincidences. Since the baseline is estimated with a single value, the thresholding method is generally effective in cases where there is little fluctuation but will not work well in situations where there are strong fluctuations (e.g. due to shared common input). To account for these fluctuations, Hatsopoulos et al. (2003) developed a pattern jitter method where jittered spike trains generate a baseline cross-correlogram that preserves slow structure in the correlogram while removing fast, transient effects such as those due to a synaptic connection. The spike jitter method is then based on testing if the peak or trough is significantly different from the local baseline estimated from the jittered spikes (see Methods for more details). In both the thresholding and the jitter methods there is no explicit model for the slow effects and fast, synaptic effects, and each cross-correlogram is treated as a separate hypothesis test. In contrast, the extended GLM uses an explicit, parametric structure for the slow and fast effects, as well as constraints based on neuron type and conduction velocity.

Since we know where the connections are in the simulations, we can compare the performance of the model-based method to the thresholding and spike jitter methods. Fig. 4A and 4B show the overall receiver operating characteristic (ROC) curves for each method, for the two simulated networks, respectively. These curves compare the true positive rate (where a true, simulated synaptic connection is detected as a connection, regardless of whether the connection was excitatory or inhibitory) and the false positive rate (where the simulated neurons were not connected, but the method detected a connection). For Simulation 1 with common inputs, the extended GLM without any network constraints (area under the curve, AUC = .96) performs better than jitter method (AUC = .94) and thresholding method (AUC = .85). With the constraints on neuron type and conduction velocity, the performance of the model-based method improves (AUC = .99). Similarly, for Simulation 2 with real presynaptic inputs, the extended GLM with constraints (AUC = .92) outperforms the model without constraints (AUC = .86), the jitter method (AUC = .87), and the threshold method (AUC = .86). The standard errors of AUC generated using bootstrap for all the methods are less than .001.

**Figure 4:**
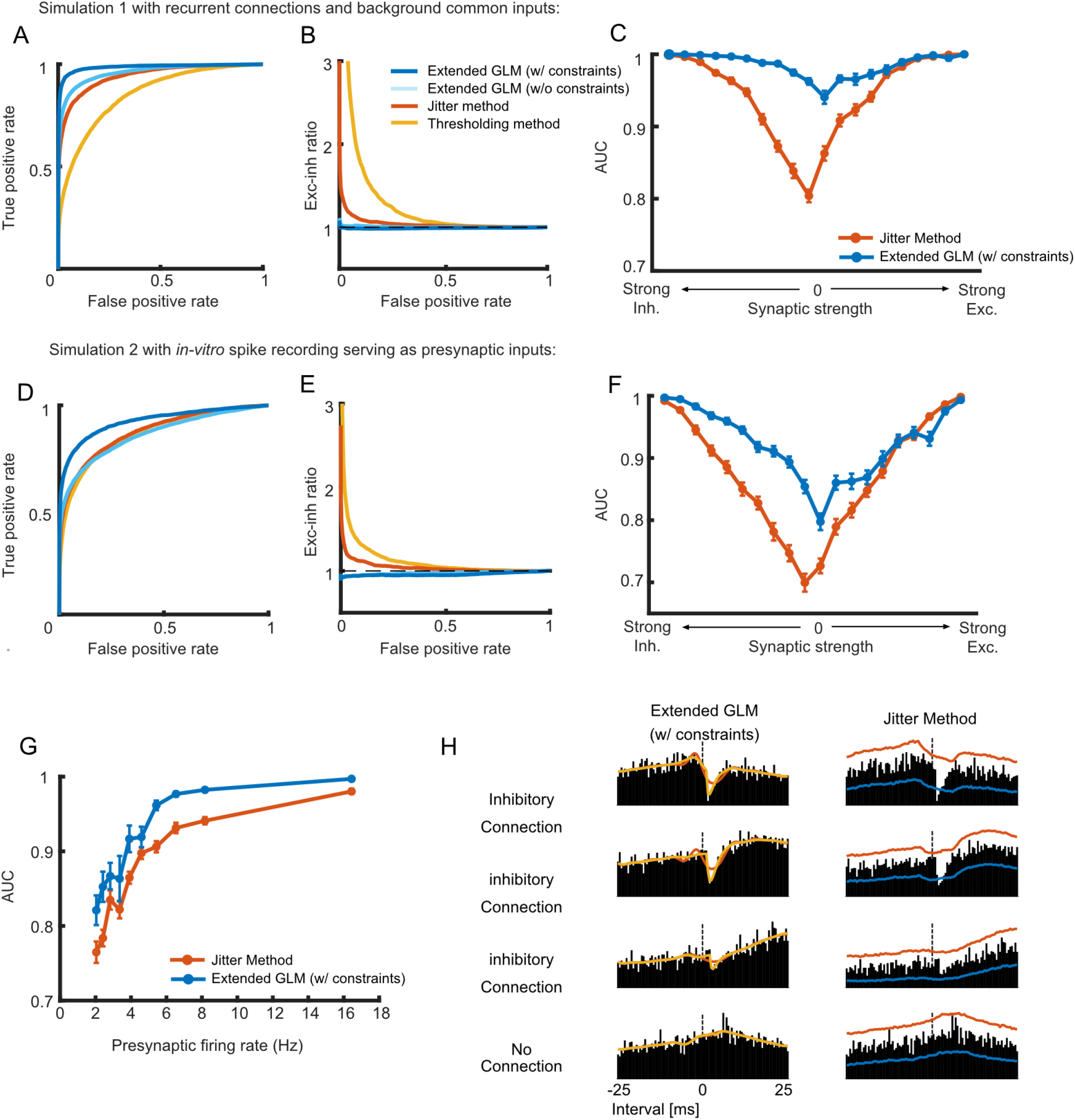
The extended GLM with constraints outperforms the jitter and thresholding methods on both of the simulations. Panel A, B and C show the results from Simulation 1 with background common inputs. Panel D, E and F show the results from Simulation 2 with real presynaptic inputs. A & D: ROC curves for the extended GLM with and without constraints, jitter method, and thresholding method. B & E: Jitter method and thresholding method are biased towards the detection of excitatory connections. The y-axis is the ratio of the excitatory true positive rate and the inhibitory true positive rate. If the method has no preference for connection type, the ratio should be 1. C & F: The extended GLM with constraints performs better than jitter method especially on weak connections. Here we divide the synaptic connections into 20 groups based on their synaptic weights and calculate AUC for each group (each group contains 5% of the connections). The error bars denote standard error (estimated using bootstrapping). G: The performance of both of the two methods is affected by the presynaptic firing rate. We divide all the presynaptic neurons into 10 groups based on their firing rates and calculate AUC for each group (each group contains 10% of the presynaptic neuron). Only results from Simulation 2 are shown, since there is a wide range of presynaptic firing rates. H: The extended GLM with constraints can better detect weak connections and rule out the false positives based on the learned structural information. The two columns show the same four cross-correlograms with the same inhibitory presynaptic neuron along with the results for the extended GLM (left) and the jitter method (right). For the model the yellow line represents the full model with inhibitory alpha function, and the red line represents the slow model. For the jitter method, the red and blue lines denote the upper and lower bounds (based on the best MCC), respectively.

Although all methods perform well above chance in detecting connections, we find that both the jitter method and thresholding method have a bias towards the detection of excitatory connections. When the decision criterion is set such that the number of false positives is small (less than ~10%) both methods detect far more excitatory connections than inhibitory connections, despite the fact that the number and strengths of excitatory and inhibitory connections were approximately balanced in the simulations. This bias may be partially due to the fact that here, for jitter method and thresholding method, we approximate the noise distribution of the correlograms using a normal distribution (z-scores), rather than using an empirical distribution. On the other hand, the extended GLM shows no preference for either excitatory or inhibitory connections for simulation 1, and a slight bias towards inhibitory connections for simulation 2 (Fig. 4B & E).

In addition to the overall performance and the performance on different cell types, we also expect the detectability of synapses to depend on the synaptic strength and the rates of the pre- and postsynaptic neurons. Here we find that, for both of the simulations, the extended GLM with constraints and the jitter method perform at a similar level for strong connections, but that the extended GLM has better detection for weak connections (Fig. 4C and F). We also find that the performance of both methods varies as a function of the firing rate of presynaptic neurons. Here the extended GLM outperforms the jitter method at all rates, but both of the methods show better performance for synaptic connections where the presynaptic firing rate is high compared to those where rate is low (Fig. 4G). By incorporating the learned network information, the extended GLM with constraints appears to better detect weak connections and rule out false positives. For example, although both the extended GLM and the jitter method can detect strong connections (Fig. 4H, top two correlograms), the jitter method has more false positives and false negatives. It may fail to detect a weak connection that does not exceed threshold (the third correlogram), or falsely detect a non-connection if there is noise that exceeds threshold (the bottom correlogram). On the other hand, if the weak connection has a sign and latency consistent with the constraints, the extended GLM can successfully detect it, and if the sign or latency are inconsistent with the constraints, the extended GLM can successfully rule this connection out (Fig. 4H).

In the above results, we set the bin size of the cross-correlograms to 0.5 ms for all the methods to tradeoff between the number of spikes within a bin and the time resolution. However, since bin size may affect performance, we also ran our model, the jitter method and the thresholding method with the bin sizes of 0.25 ms and 1 ms on the Simulation #1. We find that finer time resolution improves the performance of our model-based methods and does impair the performance of the jitter and thresholding methods. However, the best overall performance is at small bin sizes where the latency and time constant of the fast synaptic effect can be accurately estimated (Table 2).

**Table 2:** Performance (Area Under the Curve) of different methods with different bin sizes on Simulation #1.

|  | GLM<br>(w/ constraints) | GLM<br>(w/o constraints) | Jitter method | Thresholding<br>method |
| --- | --- | --- | --- | --- |
| Bin size = 0.25 ms | .9883 | .9627 | .8850 | .7837 |
| Bin size = 0.5 ms | .9852 | .9597 | .9426 | .8508 |
| Bin size = 1 ms | .9775 | .9472 | .9668 | .8772 |

The overall connection probability may also play a role in detection accuracy. Higher connection probabilities may cause more cases where pairs of neurons have spurious correlations due to common drive from a third neuron. Lower connection probabilities could also make the constraints less useful, since there will be fewer connections from which to estimate both types and distance-latency relationships. As a reference, we ran Simulation #1 with increased connection probability (20% for excitatory neurons, 80% for inhibitory). With this denser simulation the performance for all model decreased: the GLM with constraints had an AUC of .97, the GLM without constraints had an AUC of .89, the jitter method had an AUC of .90, and the thresholding method had an AUC of .78. However, the relationship among the methods remains very similar.

### Synapse detection with *in vitro* multielectrode array (MEA) data

In order to evaluate the performance of our method on real data, we apply it to spontaneous *in vitro* spike activity recorded in a mouse somatosensory cortex slice culture using a 512-electrode array (see Methods). Here we adopt two representative datasets: dataset #13 and dataset #23, and examine potential connections between neurons with >1000 spikes recorded. Before we run the model on the dataset, in order to get rid of the possible influence of spike sorting problems, we exclude the neuron pairs when there is an anomalous peak or trough right in the middle of the correlogram (<7% of pairs, see Methods for more details).

Since we don’t know the ground truth about where the synaptic connections are in the *in vitro* data, we are not able to directly measure the performance of our synapse detection methods. However, we can qualitatively assess whether or not the method gives results consistent with what we expect. We first validate whether our method can correctly classify excitatory neurons and inhibitory neurons by analyzing the shape the spike waveform of each neuron. Previous studies have shown that the excitatory neurons typically have broader spike waveforms, while the majority of inhibitory neurons have narrower spike waveforms (Barthó et al. 2004). In the two datasets used here, the neurons with broader waveforms are more likely to be classified as excitatory neurons by our model based on their putative synaptic connections, but the results for neurons with narrow waveforms are mixed (Fig. 5A). To quantify the relationship between waveform and connectivity, we fit a Gaussian mixture model with 3 components to the trough-to-peak duration and half-amplitude duration of the waveforms creating three clusters for “broad waveforms”, “narrow waveforms”, and “outliers”. After assigning each neuron to a cluster (based on the posterior probability), we analyze the consistency between the waveform shape and the neuron type given by their putative connections. From the presynaptic neurons with putative connections detected by our method, we find that 77% of the “broad-spiking” neurons are classified as putative excitatory neurons based on their connectivity, and 47% of the “narrow-spiking” neurons are classified as putative inhibitory neurons. Inhibitory, non-fast-spiking neurons with broad waveforms have been previously reported (Dehghani et al. 2016), however, excitatory neurons with narrow waveforms are unexpected. There are likely to be some cases where the extended GLM misidentifies the neuron type, however, there are also cases where neurons with narrow waveforms appear to have putative excitatory connections with typical short-latency, fast transient increases in the cross-correlograms. This may suggest that the *in vitro* recordings here contain excitatory neurons with narrow waveforms. Many single units in the MEA data here appear to be narrow due to the fact that they have triphasic waveforms. Previous work suggests that this could indicate a nearby axon (Barry 2015; Gesteland et al. 1982; Robbins et al. 2013).

**Figure 5:**
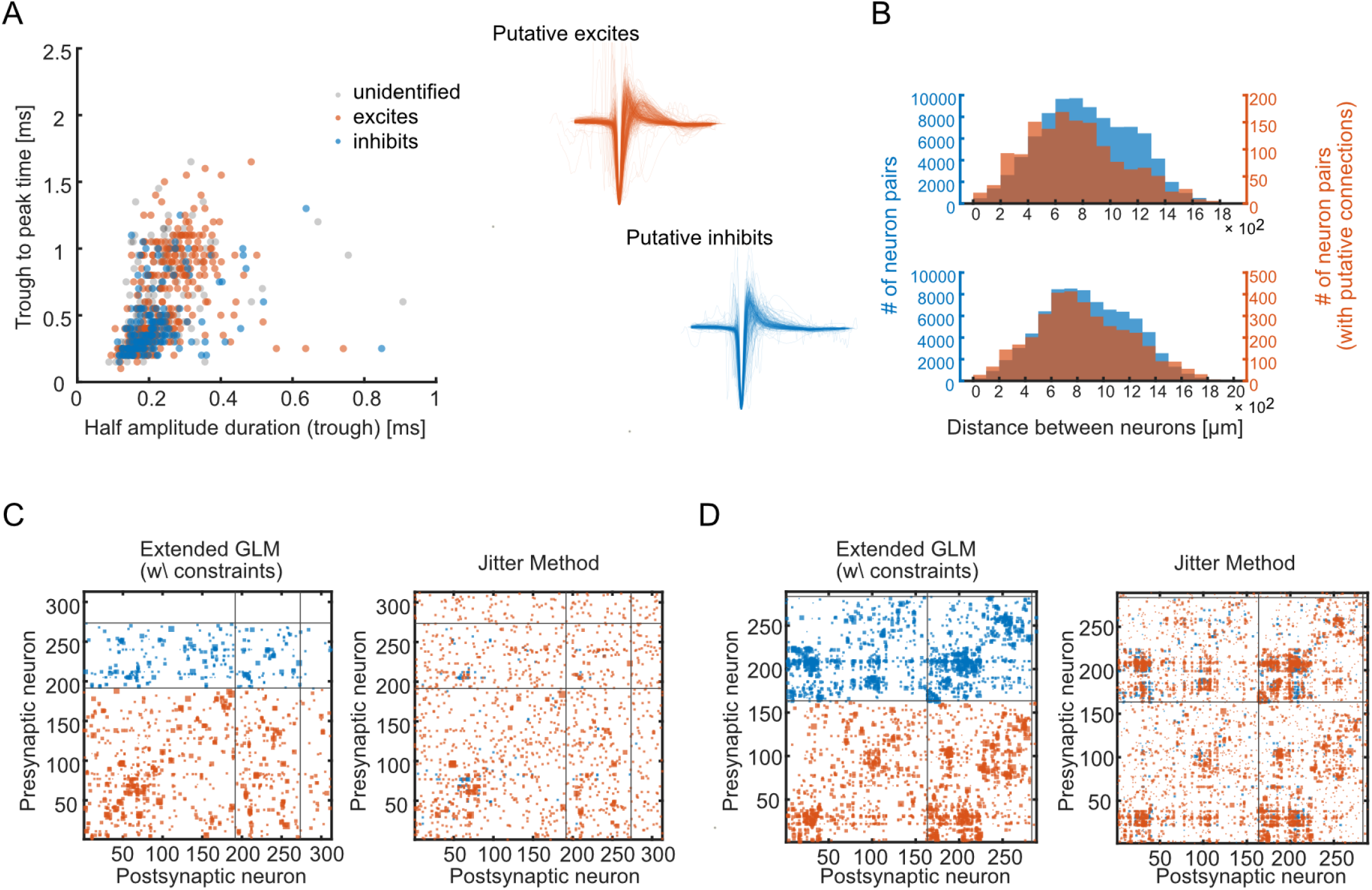
Applying the extended GLM to *in vitro* multielectrode array data. A: left: most of the neurons with wide waveforms are classified as putative excitatory neurons by our method, while the results for neurons with narrower waveforms are rather mixed. Right: The waveforms of putative excitatory neurons and inhibitory neurons. For putative inhibitory neurons, the waveforms are narrow, while for the putative excitatory neurons, there are two clusters of waveforms. B: the histograms of the distance between neurons (top: dataset #13, bottom: dataset #23). Distances for all neuron pairs are in blue, while distance for neuron pairs with putative connections are in red. C & D: Hinton plots for dataset #13 (C) and #23 (D) using the extended GLM and jitter method, respectively. The putative excitatory connections are marked in red. The putative inhibitory connections are marked in blue. Here all the neurons are sorted by the similarity of their putative connections detected by our method. Each row represents the connections from one presynaptic neuron. In each Hinton plot, the two horizontal lines separate the neurons with no putative connections, putative inhibitory neurons, and the putative excitatory neurons. The two vertical lines mark the same boundaries for postsynaptic neurons.

We then analyze the properties of the putative synaptic connections detected by our method. Here we pick the thresholds for our method based on the maximum MCC from the simulation with background fluctuations (see details in Methods). We first find that the neurons close to each other are more likely to have putative connections (Fig. 5B). The median distance between neuron pairs with putative connections is 708 μm, compared to a median distance between all the neurons of 813 μm for dataset #13. And for dataset #23, the median distance between neuron pairs with putative connections is 810 μm compared to the median distance between all the neurons 859 μm. These results are consistent with previous findings in other cortical areas that the probability of a synaptic connection decreases with distance (pyramidal cells in layer 2/3 of rat visual and somatosensory cortex: Holmgren et al. 2003; pyramidal cells in layer 5 of rat visual cortex: Song et al. 2005).

We then compare the putative connections detected by the extended GLM and the jitter method on these same datasets. As with our method, we pick the threshold for jitter method based on the maximum MCC (see details in Methods) from Simulation 1 with background fluctuations. In general, the extended GLM and jitter method detect highly distinct sets of connections (Fig. 5C and 5D). Here we sort the neurons based on the similarity of their putative connections detected by our method (using hierarchical clustering). For the Hinton plot of our method, the size of each square represents the magnitude of the estimated synaptic weight *w_i,j_* of the corresponding neuron pair. For the Hinton plot of jitter method, the size of each square represents the magnitude of the z-score of the corresponding neuron pair.

Based on the Hinton plots, we do see that the results from our method and jitter method show certain agreements on the detection of putative connections, especially on the strong connections: For dataset #13, the two methods show the same detection results (whether there is a synaptic connection or not) on 96.8% of the neuron pairs, for dataset #23, the two method show the same detection results on 94.13% of the neuron pairs. However, since the vast majority of pairs are not connected, we also use MCC to measure the similarity between the results of the two methods. The MCC between the results of the two methods is .30 (dataset #13) and .46 (dataset #23), which implies some disagreements between the results of the two methods. We find that jitter method reports more putative connections than our method (dataset #13: 2582 vs. 1423, dataset #23: 5204 vs. 3543). In addition, our method reports more putative inhibitory connections. For dataset #13, 26.4% (375 out of 1423) of the putative connections are inhibitory when using our method, while 5.9% (152 out of 2582) of the putative connections are inhibitory when using jitter method. For dataset #23, 50.7% (1797 out of 3543) of the putative connections are inhibitory when using our method, while 16.3% (846 out of 5204) of the putative connections are inhibitory when using jitter method.

We then examine to what extent the synaptic latencies of the putative connections from one presynaptic neuron increase as function of distance. For each neuron with more than 2 putative connections (450 out of 602 neurons across both datasets), we calculate the Pearson correlation coefficient *r* between the estimated synaptic latency Δ*t* and the distance between the corresponding pre and postsynaptic neuron. Fig. 6E shows the histogram of all the correlation coefficients of the two datasets, 72% of the neurons show a positive correlation between the estimated synaptic latency and distance between neurons, 36% of them are statistically significant (*p* < .05). Fig. 6A & 6B show some examples of presynaptic neurons that have many connections and are consistent with a linearly increasing latency-distance relationship (r > 0). We find both putative excitatory and putative inhibitory cases where this relationship seems to hold. In the cases where the neurons don’t obey the rule (*r* < 0, 29% of the neurons), the accuracy of the linear fit of the latency-distance relationship tends to be lower. Under the extended GLM the constraint on the synaptic latency for these ill-predicted connections (*η_Δt_*) is also weaker (although not significant, unpaired t-test:*t*(240) = −1.91, *p* = .058, *CI* = [-1.73,.03], Fig. 6F). Since the strength of the constraint in our model is partially based on how well the latency-distance relationship is fit by a linear trend, these constraints thus have a weaker influence and our method is still able to detect putative connections at unexpected latencies (Fig. 6C & 6D).

**Figure 6:**
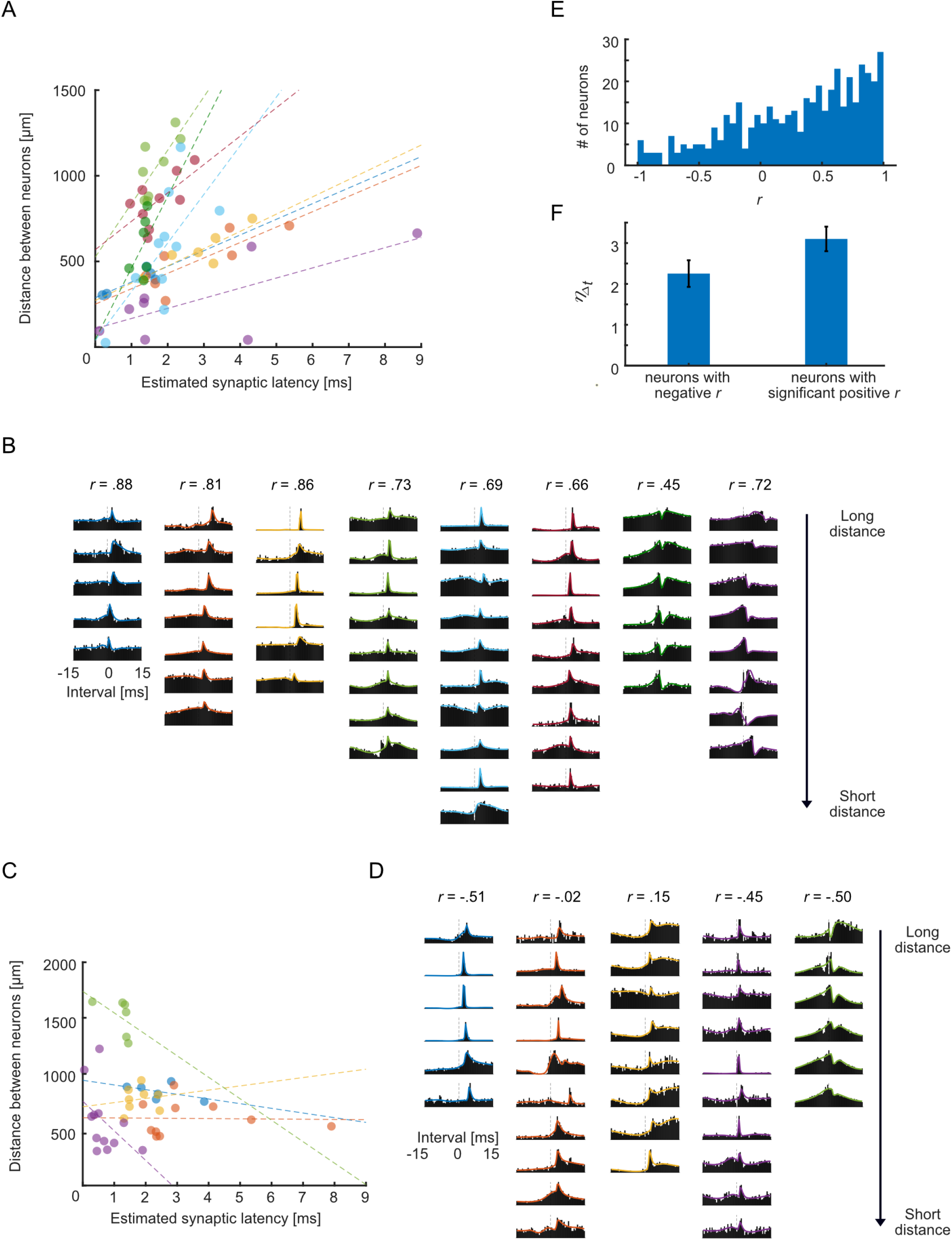
Applying the extended GLM to *in vitro* multielectrode array data. A,B, C & D: examples of neurons where the relationship between synaptic latency and distance is consistent with an increasing linear trend (panel A & B) and inconsistent with such a trend (panel C & D). Panel A & C: data points with the same color represent putative connections from the same presynaptic neuron. The dotted lines show the linear regression of the estimated synaptic latency and distance. Panel B & D : cross-correlograms for these connections with colors corresponding to the scatter plot. E: the histogram of Pearson correlation between the putative synaptic latency and distance for all the presynaptic neurons. F: mean *η_Δt_* for the neurons that don’t obey the latency rule (*r* < 0) and the neurons that obey the rule (*r* > 0 & *p* < .05).

## Discussion

Traditionally, intracellular recording represents a gold standard for characterizing synaptic connections. Detecting synaptic connections using the intracellularly recorded postsynaptic potentials and currents is straightforward and reliable (Harris et al. 2016; Song et al. 2005). However, only a relatively small number of neurons can be recorded simultaneously using intracellular recording, particularly *in vivo* (but see Pawlak et al. 2013). In recent decades, advances in multielectrode arrays have allowed the spiking of hundreds to thousands of neurons to be recorded simultaneously *in vivo* or *in vitro* with thousands of potential synapses between them (Cheung et al. 2007; Ito et al. 2014; Seeman et al. 2018; Spira and Hai 2013). Distinguishing the monosynaptic connections from the many tens of thousands of possible connections in these large-scale extracellular recordings is a difficult statistical problem. Previous methods for distinguishing putative synaptic connections and non-connections in large-scale recordings have used separate hypothesis tests on the cross-correlograms of all potentially connected neuron pairs (Hatsopoulos et al. 2003; Pastore et al. 2018; Perkel et al. 1967b). In recent work, Kobayashi et al. 2019 find that a model-based approach combining a slow background effect and a fast synaptic effect (GLMCC) provides improved performance. Here we develop an extended GLM that also incorporates two structural constraints learned from the whole network: presynaptic neuron type and the relationship between the synaptic latency and distance between pre- and postsynaptic neurons. On two simulated integrate-and-fire networks, our model outperforms previous synapse detection methods (the thresholding method and spike jitter method), especially on the weak connections. We also apply our model on *in vitro* multielectrode arrays (MEAs) data. Here our model recovers plausible connections from hundreds of neurons recorded extracellularly.

Many factors affect how likely a synaptic connection is to be detected, including the firing rates of the pre- and postsynaptic neurons, the recording time, and the synaptic strength. Here, in our simulations, we find that the model-based approach outperforms the hypothesis testing-based approaches for a wide range of firing rates and shows particular improvement for detecting weak connections. At the same time, in our simulations, the model-based methods outperform the hypothesis test-based methods at all thresholds. That is, the distributions of likelihood ratios for connections and non-connections are more distinct than the distributions of test statistics with the jitter or thresholding methods. In practice, however, when detecting putative synapses, the choice of threshold has a strong effect on how many synapses are detected and also how many false positives there are. Here, in detecting putative synapses in experimental data we apply the same optimal (MCC maximizing) threshold from the simulation. This is largely for illustration, but selecting an appropriate threshold for experimental recordings depends on the researchers’ tolerance for false positives and false negatives. Ultimately, the choice of threshold should be based on the aims of the analysis and the costs/benefits of mistakes in interpreting the underlying data.

Since we don’t know the ground truth for experimental data, it is possible that the threshold used here might be either too strict or too permissive. However, the performance of the model-based method may be somewhat more robust to the choice of threshold than the jitter and thresholding methods. In our simulations, we find that both the jitter method and thresholding method show strong biases towards detecting excitatory connections, particularly at strict thresholds with few false positives. The model-based approach, on the other hand, detects excitatory and inhibitory connections in proportion to their prevalence in the simulation at all the thresholds. The bias of the jitter method may due to the fact that we here measure test statistics assuming that spike counts follow a normal distribution. This approximation clearly does not accurately account for the fact that spike counts can only be non-negative. However, in practice we find that this type of smooth approximation has better performance at strict thresholds compared to using the empirical count distributions (using the percentile of the true counts in the jittered count distribution), which do not have smooth tails. These biases we find in the simulation results may indicate that, when we apply these methods to real data, jitter method and thresholding method may distort the observed E-I ratio if the threshold is too strict. Consistent with the simulation results, in the *in vitro* data analysis, we find that the jitter method also typically detects many more excitatory than inhibitory connections (5-13x more), while the model-based method detects putative connections with a smaller EI ratio (~3:1). Previous work has found that approximately one in five neurons is GABAergic in many neocortical areas and species (Hendry et al. 1987; Sahara et al. 2012). Although there are many factors that might influence the observed EI ratios when measuring putative synapses from spikes, the model-based approach appears to be less biased.

In the model-based approach, we learn two structural constraints from the whole network: presynaptic neuron type and the relationship between the synaptic latency and distance between pre- and postsynaptic neurons. For the presynaptic neuron type, using the simulation, we find that the model-based approach is able to successfully classify most neurons. However, when applying the method to the *in vitro* data, we compare the neuron type estimated based on putative synaptic connections with waveform shapes, and find that our results are somewhat less clear than previous findings *in vivo* (Barthó et al. 2004). Instead of two, well separated excitatory (broad waveforms) and inhibitory (narrow waveforms) clusters, we find substantial mixing of types across clusters. This may be partially due to the particulars of organotypic slice recording. Previous works have found that the waveforms in these recordings tend to be more triphasic potentially due to axonal conductance (Barry 2015; Robbins et al. 2013), and this could lead to misestimation of waveform width. New methods, such as optotagging (Lima et al. 2009) or optrodes (English et al. 2017) may offer a more reliable identification of neuron type. However, in the absence of experimental verification, it is difficult to evaluate the accuracy of cell type inferences. Additionally, although here we assume that presynaptic neurons are either exclusively excitatory or exclusively inhibitory, there is recent and growing evidence that presynaptic neurons can co-release multiple neurotransmitters (Root et al. 2014).

For the relationship between the synaptic latency and distance between pre- and postsynaptic neurons, we found that the model-based method can successfully learn linear relationships in simulation and that these constraints improve detection performance. By applying our method to the *in vitro* data, we also find that for most of the neurons, the synaptic latencies tend to increase with the distance between the pre- and postsynaptic neurons. However, there appears to be a portion of neurons that don’t show this pattern. In many cases, we may not have enough putative synaptic connections to estimate such a trend. In the cases where there are enough connections, there may not be a trend due to several other reasons. First, the locations of the somas are only approximate – based on which electrodes have the highest amplitude waveforms. Second, although here we model presynaptic conduction velocity, it’s possible that the dendritic distance constitutes a large portion of the distance. And third, the straight-line distance between somas may not be the same as the trajectory of the axons/dendrites, especially when the neurons are sampled from different barrel fields or different cortical layers. Although previous theoretical work on the minimum wiring length principle might suggest the conduction distance between two neurons can be well approximated with straight-line (Chklovskii et al. 2002; Koulakov and Chklovskii 2001), there are clearly many sources of uncertainty when estimating conduction velocity here. However, it is important to note that, within the extended GLM, the conduction velocity is only a soft constraint, and the strength of the constraint is related to how accurately the relationship is fit by a straight line. We are still able to detect connections even if the relationship between synaptic latency and distance is not clearly linear.

With the model-based method, we are able to learn the properties of each presynaptic neuron (type and conduction velocity) and use these properties to better detect individual synaptic connections based whether they are consistent with these properties. In the future application of this method, we could potentially include other sources of information to better estimate these properties. For instance, cell types can be classified according to: mean firing rate, the mode of the inter-spike interval distribution, burstiness, and spike asymmetry (English et al. 2017), using “center of mass” could provide an estimate of the directions of connections(Gerrard et al. 2008; Luczak et al. 2009; Takeuchi et al. 2011), and conduction velocity could also potentially be estimated using spatiotemporal electrical image generated using the spike waveforms across multiple electrodes (Li et al. 2015). In addition, the model-based approach is flexible enough that other constraints could also be incorporated. For instance, we could use constraints based on connectivity across and between brain regions or other network structure (Linderman et al. 2016). Additionally, although we have not focused on these measures here, the log likelihood ratios that we use to detect the connections could also be used to calculate p-values (using a likelihood ratio test), and confidence intervals for the estimated synaptic weights can be calculated based on the log likelihood itself. Finally, as neural recording techniques continue developing, increasing numbers of neurons can be recorded simultaneously (Stevenson and Kording 2011). These recordings have the potential to contain more monosynaptic connections per recording, and this should result in more reliable estimation of neuronal properties. Although the methods presented here are likely to be useful for large-scale detection of putative synaptic connections, modeling the cross-correlogram directly does not necessarily provide unambiguous evidence for or against the presence of a synapse. Other detection methods have used other spike statistics (Casadiego et al. 2018; Chen et al. 2011; Ito et al. 2011; Kadirvelu et al. 2017; Ladenbauer et al. 2019; Monasson and Cocco 2011; Song et al. 2013), and the shape of the cross-correlogram can be influenced by many other factors, such as the dynamics of the presynaptic neuron (Perkel et al. 1967b) and common input from unobserved neurons (Gerstein et al. 1989; Stevenson et al. 2008). Here we exclude the neuron pairs when there is a peak or trough in the cross-correlogram at t=0 to remove the potentially problematic connections, and expect our slow basis functions can act to model the common inputs if they occur on slow timescales (~10 ms). There are, however, examples of systems where common input on fast timescales does occur, see (Diba et al. 2014; Swadlow et al. 1998). These could potentially result in false positives under our framework. Additionally, although we account for some potential structure due to properties of presynaptic neurons, modeling multiple inputs to the same postsynaptic neuron will likely result in more accurate estimates of the true connectivity (Roudi et al. 2015; Volgushev et al. 2015; Zaytsev et al. 2015).

Ultimately, being able to accurately detect putative synaptic connections from large-scale extracellular recordings opens a host of neuroscientific questions. Previous work found that synaptic weights detected from spikes can have strong type-dependent structure (Barthó et al. 2004), seem to vary based on behavior (Fujisawa et al. 2008), and also have substantial short-term dynamics (English et al. 2017; Ghanbari et al. 2017). Although here we apply our method to an *in vitro* MEA dataset, this method can be applied to any datasets that contain spike trains and inter-neuron distances. Our method provides an additional tool for detecting putative synaptic connections in *in vivo* or *in vitro* large-scale recordings. As the scale of recording techniques increases, this approach may help us better understand how the properties of single neuronal connections relate to population neural activity and behavior.

## Code and MEA data availability

The MATLAB code of the extended GLM method is available at GitHub: https://github.com/NaixinRen/extended-GLM-for-synapse-detection.

The MEA *in vitro* data is available at the Collaborative Research in Computational Neuroscience (CRNCS) Data Sharing Initiative: https://crcns.org/data-sets/ssc/ssc-3/about-ssc-3

## Acknowledgements

This work was supported by NSF CAREER 1651396 to IHS and NSF Robust Intelligence grant 1513779 to JMB.

